# *Aggregatibacter actinomycetemcomitans* (*Aa*) cytolethal distending toxin (Cdt) modulates host phagocytic function

**DOI:** 10.1101/2023.05.10.540171

**Authors:** Taewan J. Kim, Bruce J. Shenker, Andrew Macelroy, Lisa Pankoski, Kathleen Boesze-Battaglia

**Author notes:** Corresponding Author; Kathleen Boesze-Battaglia, Department of Basic and Translational Sciences, University of Pennsylvania, School of Dental Medicine.

## Abstract

Cytolethal distending toxins (Cdt) are a family of toxins produced by several human pathogens which infect mucocutaneous tissue and induce inflammatory disease. Human macrophages exposed to *Aggregatibacter actinomycetemcomitans* (*Aa*) Cdt respond through canonical and non-canonical inflammasome activation to stimulate cytokine release. The inflammatory response is dependent on PI3K signaling blockade via the toxin’s phosphatidylinositol-3,4,5-triphosphate (PIP3) phosphatase activity; converting PIP3 to phosphatidylinsoitol-3,4-diphosphate (PI3,4P2) thereby depleting PIP3 pools. Phosphoinositides, also play a critical role in phagosome trafficking, serving as binding domains for effector proteins during phagosome maturation and subsequent fusion with lysosomes. We now demonstrate that *Aa*Cdt manipulates the phosphoinositide (PI) pools of phagosome membranes and alters Rab5 association. Exposure of macrophages to *Aa*Cdt slowed phagosome maturation and decreased phago-lysosome formation, thereby compromising macrophage phagocytic function. Moreover, macrophages exposed to Cdt showed decreased bactericidal capacity leading to increase in *Aggregatibacter actinomycetemcomitans* survival. Thus, Cdt may contribute to increased susceptibility to bacterial infection. These studies uncover an underexplored aspect of Cdt function and provide new insight into the virulence potential of Cdt in mediating the pathogenesis of disease caused by Cdt-producing organisms such as *Aa*.

## INTRODUCTION

Periodontitis is a chronic inflammatory disorder due to host response to specific oral microbiota [1, 2]. In the absence of therapeutic intervention the periodontium (consisting of tooth/teeth, alveolar bone, periodontal ligament and connective tissue attachment) can be damaged and leads to tooth loss [1, 3]. Stage 3 or 4 and Grade C with a molar-incisor pattern [4], formerly known as Localized aggressive periodontitis (LAP), is one of the most severe forms of periodontitis. *A. actinomycetemcomitans* (*Aa*), is implicated in the etiology and pathogenesis of LAP as it serves as an early colonizer that facilitates the transition from health to disease. Presumably, *Aa* creates conditions that favor colonization by other organisms; this may include suppression of the host immune system and damage to the periodontium [5-7]. Among the virulence factors produced by *Aa*, current studies suggest that the cytolethal distending toxin (Cdt) contributes to this altered environment.

Cdt is produced by over 30 γ and ε-Proteobacteria that are human and/or animal pathogens and colonize mucocutaneous tissue such as oral, gastrointestinal, urinary, and respiratory tracts [8, 9]. Such Cdt-producing bacteria induce disease in these mucocutaneous niches characterized by sustained infection and inflammation. Our recent studies demonstrate that Cdts from *Aa, Haemophilus ducreyi* (HdCdt) and *Campylobacter jejuni* (CjCdt) exhibit potent PIP3 phosphatase activity. Moreover, lymphocytes treated with these Cdts exhibit PI-3K signaling blockade: reduced levels of pAkt and pGSK3β [10]. Previous studies have suggested a novel mechanism of action in which, the active subunit, CdtB, acts as a phosphatidylinositol-3,4,5-triphosphate (PIP3) phosphatase [11]. Following internalization, CdtB converts PIP3 to phosphatidylinsoitol-3,4-diphosphate (PI3,4P2) leading to depletions in the PIP3 poolsand increases in PI3,4P2. A shift in PIP3 levels not only modulates Akt-GSK3β signaling [10, 12, 13] but also has critical implications for phagocytic function of macrophages that may lead to alterations in phagosome maturation and in turn anti-microbicidal activity [14, 15].

Phosphoinositides (PIs) play an essential role in the trafficking of endosomes and phagosomes by serving as spatio-temporal signposts that direct maturation of phagosomes for content degradation [14, 16]. Phagosomes are dynamic structures that interact with endosomes in a process involving acquisition and release of membrane and luminal components as the phagosome matures to a phago-lysosome [16]. This maturation process is controlled by recruitment of proteins, such as Rab5 and Rab7, which are regulated by PI distribution [14, 17]. Rab5 to Rab7 conversion drives transition of early phagosomes to late phagolysosomes [14, 17]. PI interconversion can be disrupted either from naturally occurring mutations in PI converting enzymes (phosphatases and kinases) or by experimental manipulation of expression of these enzymes [16, 18, 19]. Such enzymatic defects alter PI distribution, disrupt vesicular transport and are the underlying cause of disease such as oculocerebrorenal syndrome of Lowe, neurological disorders, cancer, and numerous intercellular pathogen related disorders such as tuberculosis, legionellosis, typhoid, and listeriosis [20, 21]. The alteration in PI pools and the resulting impairment of phago-lysosome formation can significantly adversely affect macrophage host defense function. In this study, we tested that hypothesis that Cdt, via its PI phosphatase activity, hijacks phagocyte maturation thereby creating an intracellular niche that supports *Aa* survival.

## MATERIALS AND METHODS

### Reagents and Antibodies

The following antibodies were utilized for western blot and immunofluorescence studies. Mouse anti-PI3,4P2 mAb and mouse anti-PI3,4,5P3 mAb conjugated with FITC from Echelon Bioscience (Salt Lake City, UT). Rabbit anti-EEA1 mAb from Cell Signaling Technology (Danvers, MA). Goat anti-rabbit Ig-HRP conjugate, goat anti-mouse Ig-HRP conjugate, and goat anti-mouse pAb conjugated with Alexa Fluro 488 from Invitrogen (Carlsbad, CA). Rabbit anti-LAMP1 pAb, and rabbit anti-Rab5 pAb from Abcam (Cambridge, England). Mouse anti-Rab7 mAb from Sigma Aldrich (St. Louis, MO). For phagocytosis and phagosome maturation assays, pHrodo™ Red *E. coli* BioParticles™ Conjugate and DQ™-BSA Red (used in 96 well plate assay) and DQ™-BSA -Green (used for live cell assay) was purchased from Invitrogen.

| Antibody (Host) | Source; Catalog # | Dilution (application) |
| --- | --- | --- |
| Anti-PI3,4P2 (mouse) | Echelon; Z-P034 | 1:100 (Immuno) |
| Anti-PI3,4,5P3 (mouse) | Echelon; Z-G345 | 1:50 (Immuno) |
| Anti-EEA1 (rabbit) | Cell Signaling; C45B10 | 1:100 (Immuno); 1:1,000 (Western) |
| Anti-LAMP1 (rabbit) | Abcam; ab24170 | 1:100 (Immuno); 1:1,000 (Western) |
| Anti-Rab5 (rabbit) | Abcam; ab18211 | 1:100 (Immuno); 1:1,000 (Western) |
| Anti-Rab7 (mouse) | Sigma-Aldrich; R8779 | 1:100 (Immuno); 1:1,000 (Western) |

### Cell Culture

The human acute monocytic leukemia cell line, THP-1, was obtained from ATCC (Manassas, VA); cells was maintained in RPMI1640-containing 10% FBS, 1 mM sodium pyruvate, 20μM 2-mercaptoethanol and 2% penicillin-streptomycin at 37°C with 5% CO_2_ in a humidified incubator. THP-1 cells were differentiated into macrophages by incubating cells in the presence of 50ng/ml PMA for 48 hours at which time the cells were washed and incubated an additional 24 h in medium prior to use [13].

### Expression and purification of Cdt, CdtB mutants and Cdt holotoxin

Construction and expression of the plasmid containing the cdt genes for the holotoxin (pUCAacdtABChis) have previously been reported [22]. The plasmid was constructed so that cdt genes were under control of the lac promoter and transformed into *E. coli* DH5α. Cultures of transformed *E. coli* were grown in 1L LB broth and induced with 0.1 mM of isopropyl β-D-1-thiogalactopyranoside for 2 h; bacterial cells were harvested, washed, and resuspended in 50 mM of Tris (pH 8.0). The cells were frozen overnight, thawed, and sonicated. The histidine-tagged peptide holotoxin was isolated by nickel affinity chromatography as previously described [23].

### Measurement of cellular PI(3,4,5)P3, PI(4,5)P2, and PI(3,4)P2 content

Cells (1×10^6^ cells/well) were incubated with Cdts or media for 240 min. Replicate cultures (0.5–1×10^7^cells) were pooled and harvested. Lipids were extracted as described. [13]. The cell pellet was treated with cold 0.5 TCA for 5 min, centrifuged and washed twice with 5% TCA containing 1 mM EDTA. Neutral lipids were extracted twice with methanol:chloroform (2:1) at room temperature. Acidic lipids were extracted with methanol:chloroform:12M HCl (80:40:1) for 15 min at room temperature; the samples were centrifuged for 5 min and the supernatant recovered. The supernatant was then treated with 0.75 ml chloroform and 0.1 M HCl and centrifuged to separate organic and aqueous phases; the organic phase was collected and dried. The dried lipids were resuspended in 120μl 50 mM Hepes buffer (pH 7.4) containing 150 mM NaCl and 1.5% sodium cholate and left overnight at 4°C. PI(4,5)P3, PI(3,4)P3, and PI(3,4,5)P3 levels were then determined using commercially available competitive ELISA according to the manufacturer’s directions (Echelon).

### Confocal Microscopy

#### Phosphoinositide immunofluorescence

Cells were treated with Cdts and fixed with 2% PFA and permeabilized with 0.05% saponin. After blocking with 10% goat serum (SouthernBiotech; Birmingham, AL), samples were incubated with mouse anti-PI(3,4)P2 mAB or mouse anti-PI(3,4,5)P3 mAB conjugated with FITC for 1 hour at 37°C [24]. TBS with 1% goat serum was used for washing. For PI(3,4)P2 secondary antibody (goat anti-mouse) conjugated to Alexa Fluor 488 was used while no secondary antibody was used for PI(3,4,5)P3 which was directly conjugated with FITC. Hoechst 33258 (AnaSpec Inc; Freemont, CA) for all the sets and incubate at 37°C for 1 hour [25]. Cells were washed with Tris-buffered saline (TBS) and imaged within 24 hours.

#### Latex bead – protein association

Cells were treated with Cdts and human serum type AB (Atlanta biologicals S40110; Kolkata, India) opsonized red (λEx 580nm/λEm 605nm) 1μm diameter latex beads (Invitrogen F13083) were fed to stimulate phagocytosis. At time points (indicated in figure legends) cells were fixed with 4% PFA and blocked in 0.1% goat serum and 0.1% saponin and incubated with rabbit anti-Rab5, rabbit anti-EEA1, rabbit anti-LAMP1 or mouse anti-Rab7 for 18 hours at 4°C. After primary antibody incubation, cells were stained with Hoechst 33258 (AnaSpec Inc; Freemont, CA) and anti-rabbit or anti-mouse secondary antibody conjugated with Alexa-flour 488 for 1 hour at 37°C, washed with PBS, and imaged within 24 hours [26].

#### Confocal Imaging

Samples were treated and fixed in 35mm glass bottom dishes (MatTek; Ashland, MA) and images captured with a Nikon A1R laser scanning confocal microscope with a PLAN APO VC 60×water (NA 1.2) or 100×oil (NA 1.45) objective at room temperature. Image z-stacks were acquired at an interval of 0.1μm (50 focal planes/image stack, 5μm). Data were analyzed using Nikon Elements AR 4.30.01 software by maximum intensity projection and standard LUT adjustment [27]. For latex bead-PI(3,4)P2 association and latex bead–protein association studies, we measured fluorescence intensity around the latex bead (1μm diameter). The area of analysis was confined to a region centered around the bead in the Z axis and intensity measured within a 2μm x 2μm area of association [28-30].

#### Live cell imaging

THP-1 macrophages were differentiated on 35mm glass bottom dishes and treated with 500ng/ml CdtB^WT^ or media change (untreated control) for 4 hours. pHrodo™ Red *E. coli* BioParticles™ conjugate or DQ™ Green BSA was added for 30min at 4°C and moved to Cytoseal mounting medium (Electron Microscopy Sciences, Hatfield, PA) set at 37°C with 5% CO_2_. Images were captured with a Nikon A1R laser scanning confocal microscope with a PLAN APO VC 100×oil (NA 1.2) or PLAN APO VC 20×air (NA 0.75) every 30 minutes [31].

### Western Blot Analysis

Cells were treated with Cdts and solubilized in 20mM Tris-HCl buffer (pH7.5) containing 150 mM NaCl, 1 mM EDTA, 1% NP-40, 1% sodium deoxycholate, and protease inhibitor cocktail (ThermoFisher Scientific; Waltham, MA). Samples (15μg) were separated on 12% SDS-PAGE and then transferred to PVDF membranes. The membrane was blocked with BLOTTO and then incubated with primary antibodies overnight (18hrs) at 4°C [26]. Membranes were washed and incubated with secondary antibodies conjugated to horseradish peroxidase [26]. Western blots were developed using chemiluminescence and analyzed by digital densitometry (LiCor Biosciences; Lincoln, NE) [32]. Each protein was normalized to either actin or GAPDH.

### Assessment of phagocytosis and phagosome maturation activity in THP-1 cells

#### Phagocytosis Assay

THP-1 macrophages were differentiated on 96-wells plate. Cells were treated with Cdts with different concentrations as indicated for 4 hours and pHrodo™ Red *E. coli* BioParticles™ Conjugate was applied to measure the phagocytic response. Fluorescence, indication of decrease in pH, was measured at 180 minutes (544nm excitation and 590nm emission) with Multiskan FC (Thermo Scientific) [31].

#### Phagosome Maturation Assay

THP-1 macrophages were differentiated on 96-wells plate. Cells were treated with Cdts with different concentrations as indicated for 4 hours and DQ™ Red BSA was applied to measure the phagosome maturation. Fluorescence, indication of phagolysosome formation, was measured at 180 minutes (590nm excitation and 620nm emission) with Multiskan FC (Thermo Scientific) [33, 34].

### Bacteria and Growth Curve

*A*.*actinomycetemcomitans* strains, D7S-SA (wild type *Aa*) and D7S-SA CHE001 (Cdt deficient *Aa* mutant), were obtained as described by Nalbant, A., et al [35]. Each strain was plated on AAGM agar which consisted of 20g of BBL trypticase soy agar (BD; Sparks, MD); 3g of yeast extract (ThermoFisher) supplemented with 0.4% sodium bicarbonate and 0.8% dextrose [36]. After bacteria were grown on plates for 24 or 48 hours in the incubator with 10% CO_2_ at 37°C, they were inoculated in 10ml of AAGM broth until OD_600_ close to 0.2. Bacteria were plated from different dilutions at various time points on AAGM agar plates and incubated for 24 hours in the incubator with 10% CO2 at 37°C [35]. OD_600_ was measured using DU 650 Spectrophotometer (Beckman Coulter, Indianapolis IN) for each time point.

### Bacterial survival in THP-1 macrophages

THP-1 macrophages (10^6^cells/well) were differentiated on 12 well plates and incubated in the presence of medium only or with Cdt for 240 min. *Aa* strains were cultured until log phase (OD_600_ of 0.4 to 1) and used to inoculate with Multiplicity of Infection (MOI) (eukaryotic cells: bacteria) of 1:10 for 14 hours at 37°C with 5% CO_2_. Cells were then washed with PBS three times and treated with gentamicin (50μg/ml) to wash away and kill extracellular bacteria [37, 38]. Soy broth with 1% saponin (500μl/well) was added and the mixture incubated at 37°C with 10% CO_2_ for 15 min to lyse macrophages [39]. Lysates were plated (100μl/plate) on AAGM agar plates and incubated for 1 to 2 days at 37°C with 10% CO_2_. The number of survived bacteria was determined by the number of colonies forming unit (CFU).

Intracellular *Aa* levels were estimated by comparing the number of CFUs to an estimate of bacterial biomass using real-time PCR by adapting Ando-Suguimoto et al. [38]. Total DNA was obtained from the lysed inoculated THP-1 macrophages using DNeasy Blood & Tissue Kit (Qiagen, Germany). The amplification reaction was performed in a 20μl final volume containing 1μl of template DNA, 0.5μL each *A. actinomycetemcomitans* species-specific 16SrRNA primers (25pmol/μl) <forward: 5°GGCACGTAGGCGGACCTT3° and reverse: 5°ACCAGGGCTAAAGCCCAATC3°> (Invitrogen, Waltham MA) and 10μL of PowerUp™ SYBR™ Green Master Mix (Applied Biosystems, Waltham MA) [38]. Reactions were performed using an initial denaturation step at 50°C for 2mins to 95°C for 2mins followed by 40 cycles at 95°C/15sec, 58.8°C/15sec, 72°C/1min, and then followed by two steps at 95°C /15sec and 65°C /1 min and a final step at 60–95°C (ramp) for 15sec in a QuantStudioTM 3 Rea-Time PCR Instrument (Applied Biosystem, Waltham MA). Normalized CFU counts were determined using the following equation:

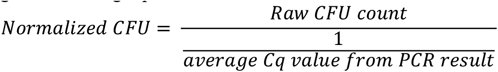

## RESULTS

Cdt treated macrophages exhibit perturbations in intracellular PI pools consistent with the CdtB subunit’s function as a PIP3 phosphatase [13]. Cells were exposed to either Cdt holotoxin containing the wildtype CdtB submit (Cdt^WT^) or Cdt holotoxin containing the phosphatase deficient CdtB subunit (Cdt^R117A^). Macrophages treated with Cdt^WT^ exhibited a two-fold increase in PI(3,4)P2 levels with a corresponding decrease in PI(3,4,5)P3 levels (Figure1A). As expected due to CdtB’s enzymatic 5’-phosphatase specificity, no change in PI(4,5)P2 was observed (Figure 1A). Furthermore, no change in PI(3,4)P2 or PI(3,4,5)P3 levels was observed in cells treated with Cdt^R117A^ (Figure 1A). The increase in PI(3,4)P2 observed with Cdt^WT^ appeared to be associated with intracellular and plasma membrane regions (indicated by white arrows, Figure 1B); semi-quantitative analyses indicated a 50% increase in PI(3,4)P2 association within these regions. No increase in PI(3,4)P2 levels were observed in cells treated with CdtB^R117A^. Figure 1C, showed distinct PI(3,4)P2 domains (white arrows) with CdtB^WT^ concentrations of 125ng/ml and higher. There was a ∼35% increase in PI(3,4)P2 with 125ng/ml and an almost 80% increase when cells were treated with 500ng/ml Cdt^WT^ [11].

**Figure 1.**
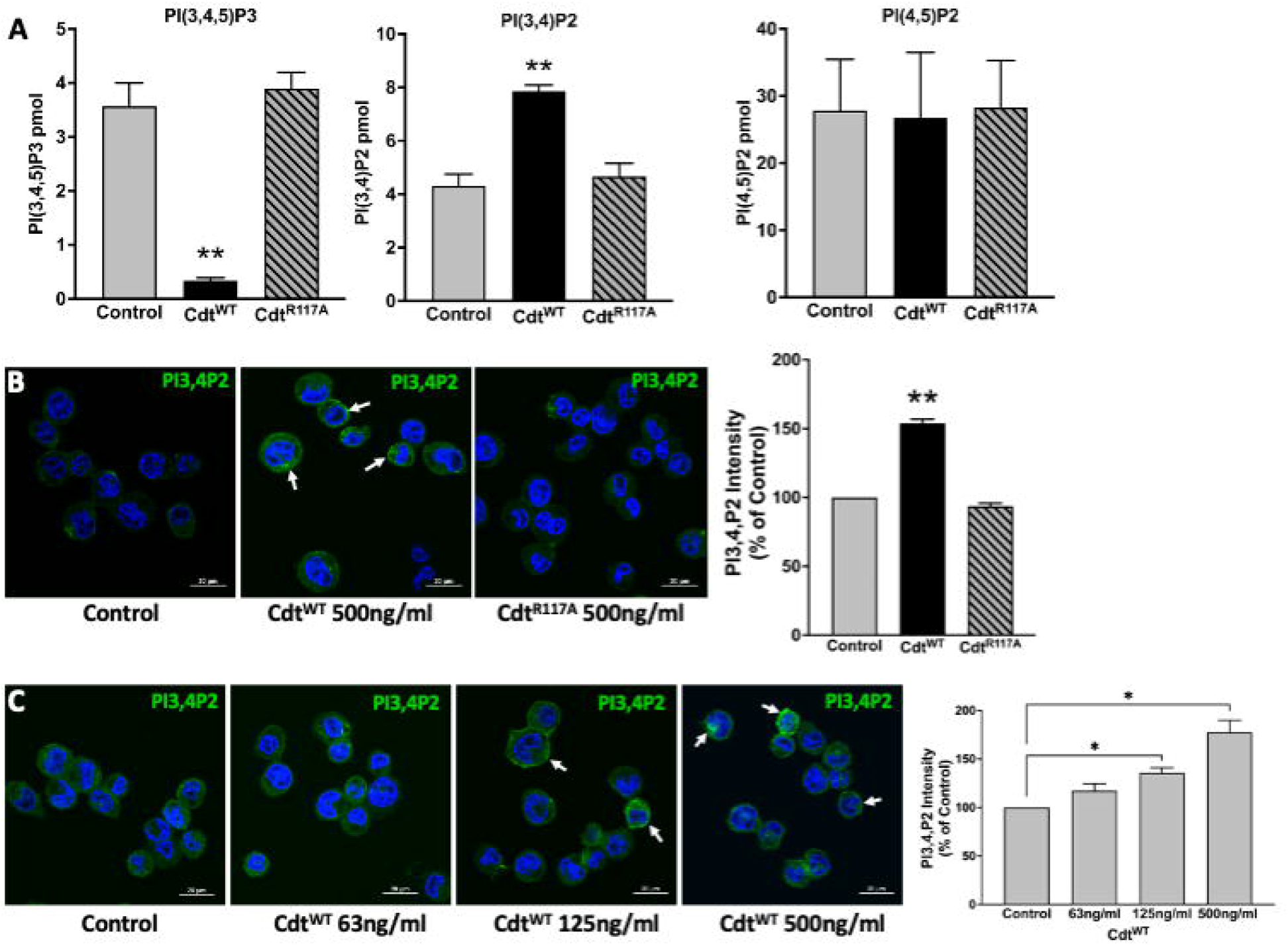
Specificity of *Aa* CdtB phosphatase activity. A. Intracellular PI pools are modulated by CdtB. PI(3,4,5)P3, PI(3,4)P2 and PI(4,5)P2, levels were measured by ELISA in extracts prepared from THP1 macrophages treated with Cdt^WT^ (500 ng/ml) or Cdt^R117A^ (500 ng/ml) as in Methods. Data represent mean +/- STDEV (n=5) and compared using Student’s t-test, **, p<0.05. B. Visualization and localization of PI3,4P2 in THP1 macrophages treated with Cdt^WT^ (500 ng/ml) or Cdt^R117A^ (500 ng/ml). PI3,4P2 (Green) as indicated by white arrows, Nucleus (Blue). Quantification of PI(3,4)P2 florescence intensity presented as percent relative to control (untreated). Data represent mean +/- STDEV (five fields with average of 12 cells/field) and compared using Student’s t-test, **, p<0.05. C. *Aa* Cdt modulates PI3,4P2 intracellular pools in dose dependent manner. THP1 macrophages were treated with Cdt^WT^ (500 ng/ml, 125ng/ml, 63ng/ml) and control (untreated) as in Methods. Visualization of PI3,4P2 level in dose dependent manner, PI3,4P2 (green, as indicated by white arrows), Nucleus (Blue). Quantification of PI(3,4)P2 fluorescence intensity presented as percent relative to the control (untreated). Results are expressed as percent of control (untreated), mean +/- STDEV (fivefields with average of 12 cells/field) and compared using Student’s t-test. *, p-value <0.05 vs. untreated control.

We next assessed the ability of Cdt to modulate macrophage phagocytic function with an opsonized pHrodo™ red E. coli BioParticles™ conjugate using live cell imaging. In preliminary experiments we determined that 10μg/ml pHrodo™ was ideal for imaging studies, (data not shown). The red pH-sensitive fluorogenic dye contributing to pHrodo™ intensity was followed for 180min, during which time intensity in control (untreated) cells started to dramatically increase at 90 min (4×10^6^ to 7×10^6^ intensity) and again at 180min (1.7×10^7^ intensity,). In contrast, when cells were pre-treated with Cdt^WT^ (500ng/ml) the magnitude of change in pHrodo fluorescence intensity was comparatively suppressed with minimum of 1.6×10^6^ intensity to maximum of 2.7×10^6^ intensity, at 180 min. The maximum intensity observed with Cdt^WT^ treatment was less than 20% of that observed in control at 180min (Figure 2A), suggesting alterations in macrophage phagocytic function. Subsequently, using a 96 well plate format, macrophages were pre-treated with Cdt^WT^ or Cdt^R117A^, at concentrations of 125-500ng/ml, prior to the addition of pHrodo™. Macrophage phagocytic function, represented by pHrodo™ fluorescence intensity, decreased by over 40% in cells pretreated with 250ng/ml Cdt^WT^ and by over 50% with 500ng/ml Cdt^WT^ (Figure 2B). There was no change in pHrodo intensity compared to controls when cells were pretreated with Cdt^R117A^ at any concentration studied (Figure 2B).

**Figure 2.**
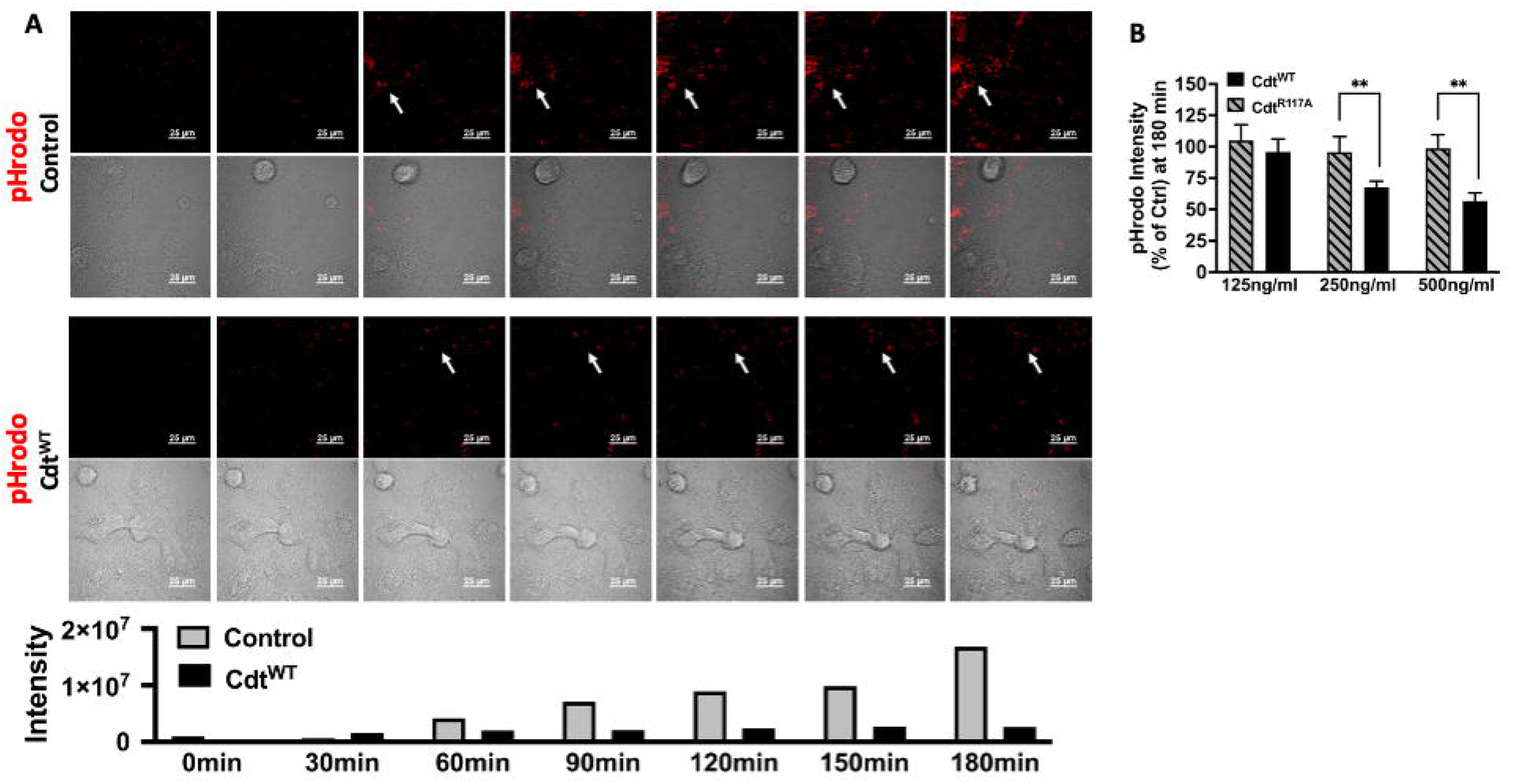
Cdt Phosphatase activity modulates host phagocytic activity. A. Time lapse imaging of pHrodo™ in THP1 macrophages pre-treated with Cdt^WT^ or control (untreated) for 4 hours. pHrodo™ fluorescence is indicated by white arrows every 30 minutes over 3 hours (Top panel: Control (untreated), Bottom panel: Cdt^WT^ treated). Intensity measurements are plotted over the experimental time course in the corresponding graph. B. pHrodo™ intensity at 180 minutes, using a 96 well plate format is shown as a function of increasing concentrations of Cdt^WT^ or Cdt^R117A^ as percent of control (untreated) as described in Methods. Data represent mean +/- STDEV (5 wells per individual experiment with 1×10^6^ cells/well) and compared using Student’s t-test. **, p-value <0.005 Cdt^WT^ vs Cdt^R117A^.

The pHrodo™ studies suggested that Cdt^WT^ may modulate phagosome maturation, a process in which phosphoinositides target maturation effector proteins. We went on to determine if there was a spatio-temporal relationship between phosphoinositide pools and ingested latex beads. Using red latex beads (1μm diameter) and the time course and Cdt^WT^ concentration (500ng/ml) established above with pHrodo™ we analyzed the area around the phagocytosed beads using multi-fluor confocal imaging. For latex bead-PI(3,4)P2 association, the area of analysis was confined to a region centered around the bead in the Z axis with fluorescence intensity measured within a 2μm x 2μm area of association. When opsonized latex beads were used as the phagocytic cargo, a dominant PI(3,4)P2 pool was observed associated with the phagocytosed bead in the presence of Cdt^WT^ (Figure 3A). There was no detectable PI(3,4)P2-latex bead association when macrophages were pretreated with Cdt^R117A^ or in untreated controls (Figure 3B and C).

**Figure 3.**
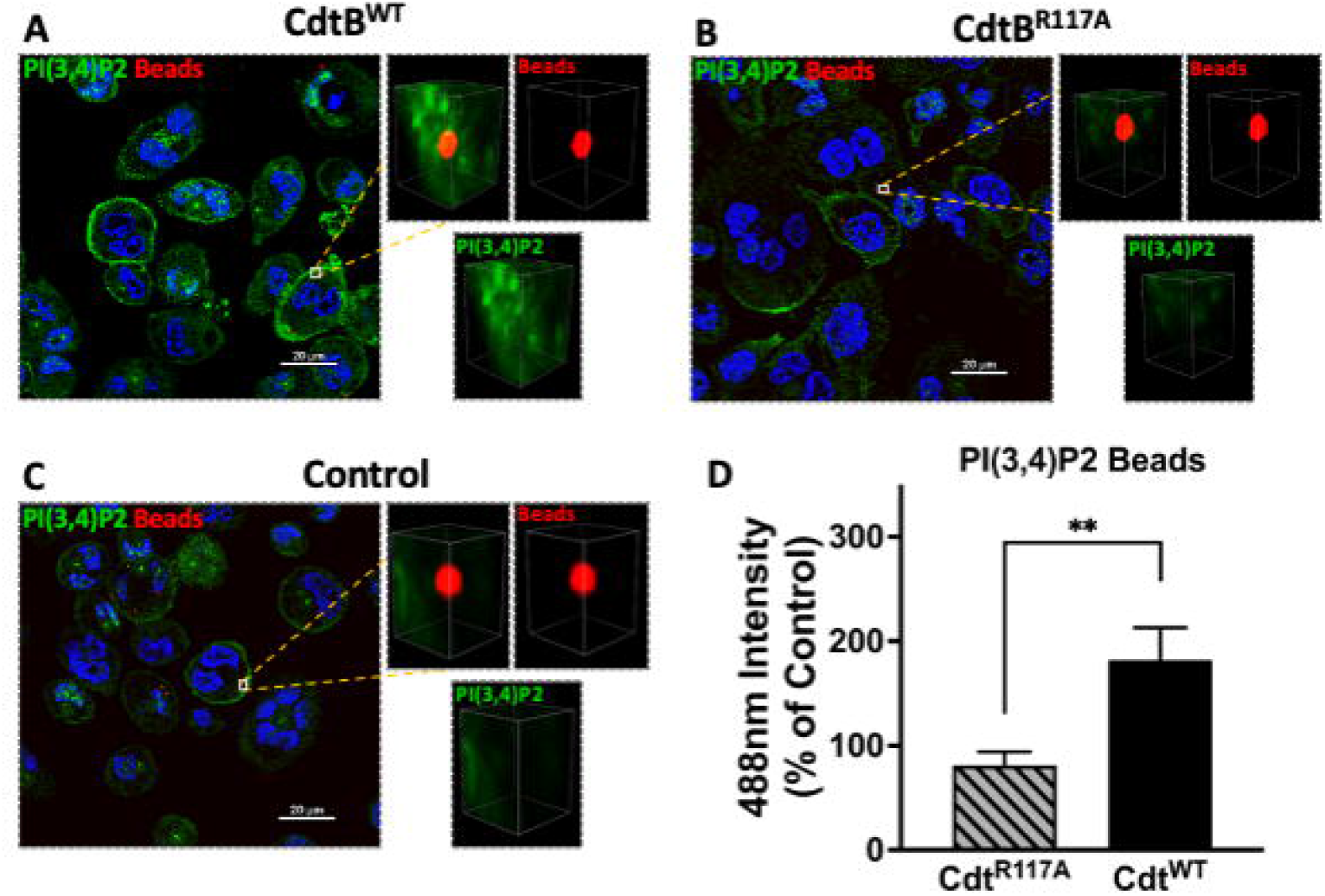
PI(3,4)P2 phagosome association increases with Cdt treatment. THP1 macrophages treated for 4 hours with Cdt^WT^, Cdt^R117A^, or control (untreated) prior to synchronized uptake of opsonized beads were fixed and stained for PI(3,4)P2 as in Methods. Green: PI(3,4)P2, Blue: nucleus, and Red: opsonized beads. Data represent average +/- STDEV of PI(3,4)P2 intensity in 2×2μm area (square) around the bead (10 beads for each for the conditions from 5 random fields) as a percent to control and compared using Student’s t-test. **, p<0.05, vs Cdt^R117A^.

We next sought to determine if Cdt modulated phagosome association with effector proteins, EEA1, and Rab5. In these latex bead–protein association studies, we measured fluorescence intensity around the 1μm diameter latex bead. The area of analysis was confined to a region centered around the bead as in the latex bead-PI(3,4)P2 association studies above (Figure 3). Using opsonized latex beads as the phagocytic cargo, we followed the association of Rab5 and EEA1 with this cargo in the presence of Cdt^WT^ (500ng/ml) using multi-fluor confocal imaging. In macrophages treated with Cdt^WT^, we observed a decrease in Rab5 association with beads when a 2μm square area around the latex bead was analyzed for Rab5 recruitment. At 30 min the association of Rab5 with beads decreased by 28% with a further decrease to 32% at 60 min, compared to untreated control cells (Figure 4A). Rab5-latex bead association was also unchanged in cells pretreated with Cdt^R117A^ (Figure 4C) implicating CdtB phosphatase activity as decreasing dissociation of phagosome-Rab5.

**Figure 4.**
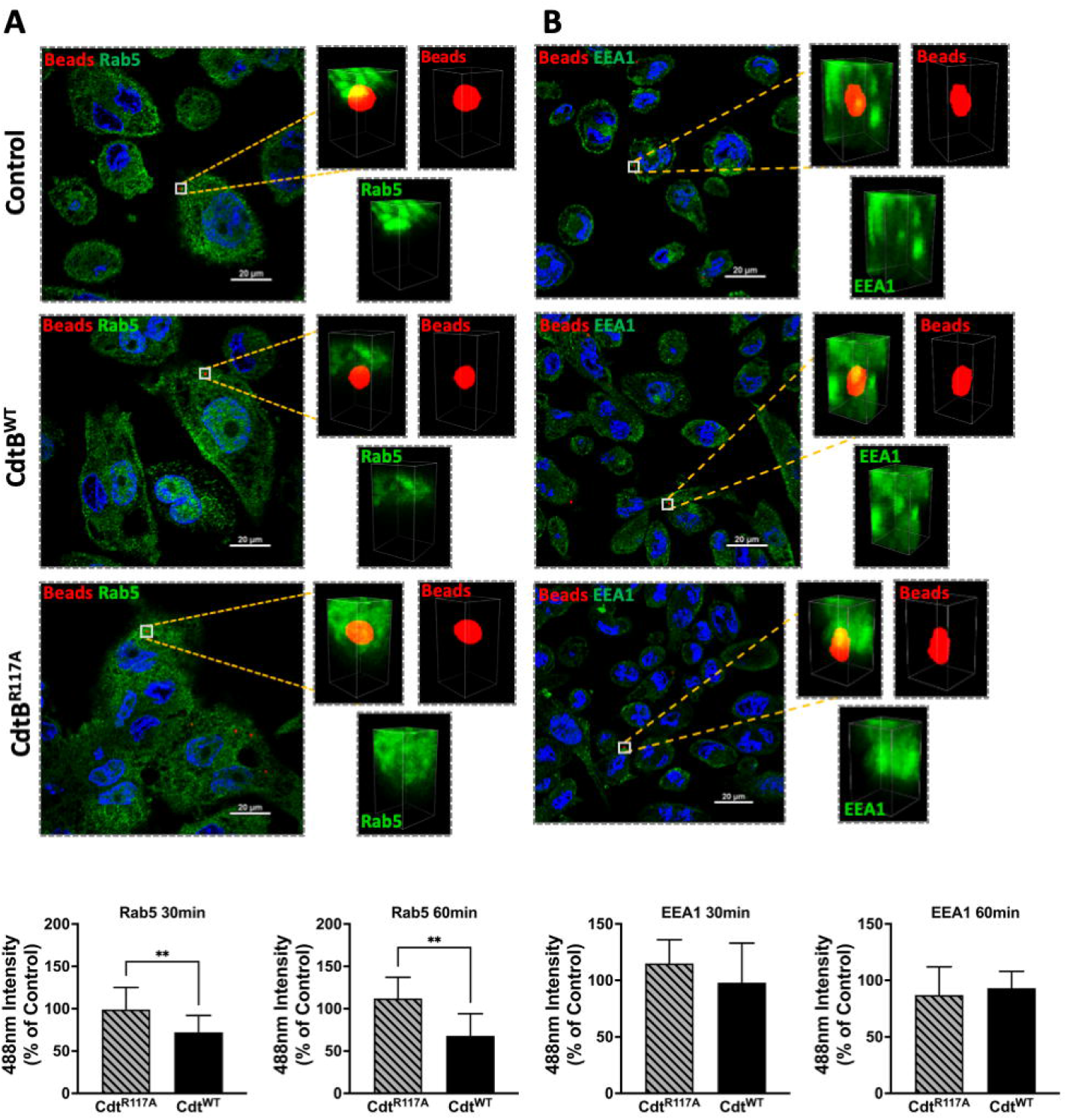
Cdt treatment increases Rab5 phagosome association. THP1 macrophages treated for 4 hours with Cdt^WT^ (500ng/ml), Cdt^R117A^ (500ng/ml) or control (untreated) prior to synchronized uptake of opsonized beads and were fixed and stained for Rab5 and EEA1 as in Methods. Green: Rab5 (Panel A) or EEA1 (Panel B), Blue: nucleus, and Red: opsonized beads. Data represent mean +/- STDEV of Rab5 (Panel A) or EEA1 (Panel B) intensity in 2×2μm area (square) around the bead (10 beads for each for the conditions from 5 different fields) as percent relative to control (untreated) and compared using Student’s t-test. **, p-value <0.05 Cdt^WT^ vs Cdt^R117A^.

Both Rab5 and EEA1 are associated with phagosomes early in the maturation process. In contrast to Rab5 association, EEA1-latex bead association was unaltered in macrophages treated with Cdt^WT^ or Cdt^R117A^ at both the 30 and 60 min time points (Figure 4B). Moreover, no change in latex-bead association with either Rab7 or LAMP1 was observed in Cdt^WT^ or Cdt^R117A^ treated cells (S. Figure 1) compared to untreated controls. Neither Cdt^WT^ nor Cdt^R117A^ (500ng/ml) treatment of macrophages altered the levels of the intracellular trafficking proteins, EEA1, Rab5, Rab7 or LAMP1, compared to untreated control (Figure 5). Collectively, these studies suggest that CdtB phosphatase activity dependent changes in phosphoinositide decreases Rab 5 association with phagosomes thereby delaying phagosome maturation.

**Figure 5.**
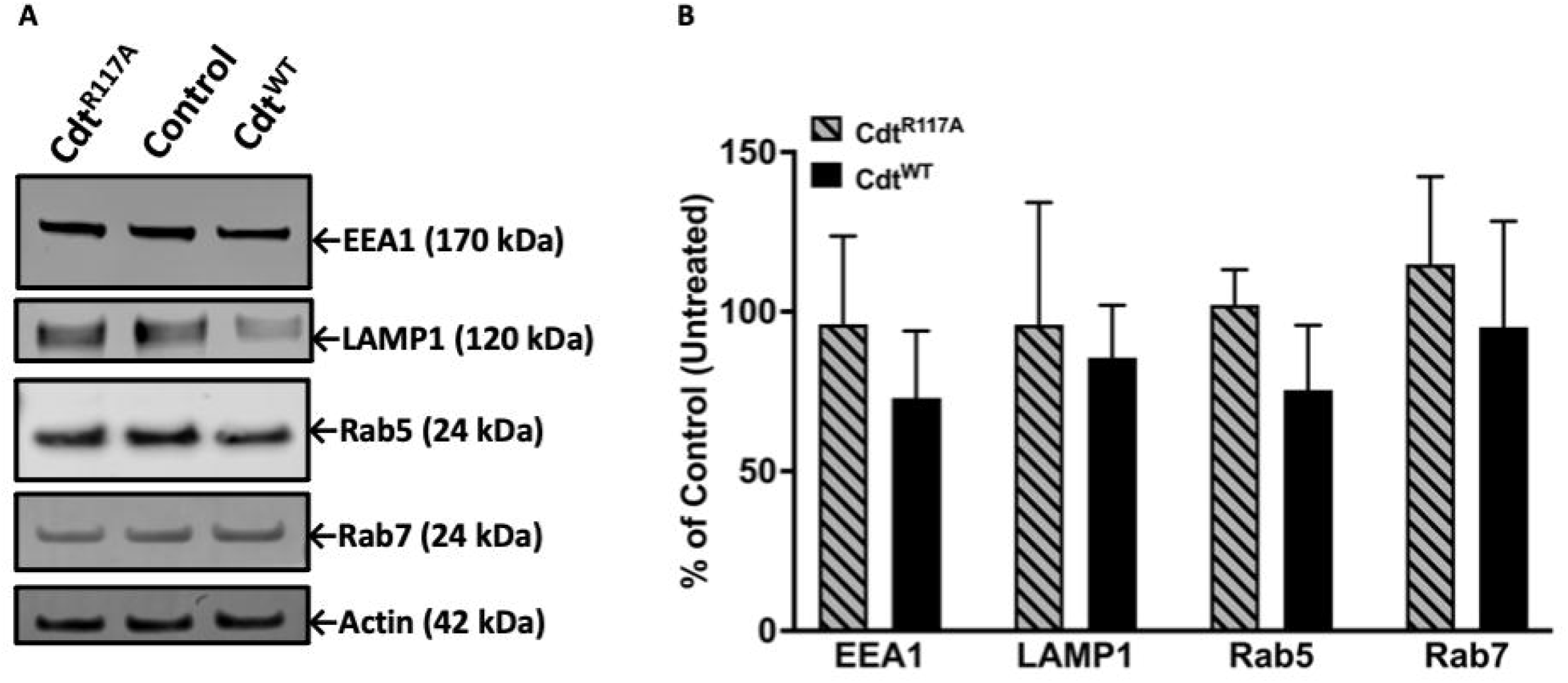
Cdt treatment does not alter levels of intracellular trafficking proteins, EEA1, Rab5 and Rab7. THP-1 macrophages treated with Cdt^WT^, Cdt^R117A^, or control (untreated). Cells were challenged with opsonized latex beads for 30 minutes and lysates collected for western blot as in Methods. Data represent mean +/- STDEV of EEA1, Rab5, and Rab7 levels (n=3) as percent of control (untreated) and compared using Student’s t-test.

The pHrodo™ studies suggested decreased phagocytic capacity but they did not specifically focus on later steps in phagosome maturation. To determine if CdtB phosphatase activity modulates phagolysosome formation, we followed DQ™-BSA fluorescence in macrophages treated with Cdt^WT^ or Cdt^R117A^. Cleavage of the self-quenched Green DQ™-BSA protease substrate in an acidic compartment generates a highly fluorescent product [34]. DQ™-BSA intensity was followed for 180min, the control (untreated) cells reached maximum intensity (5.5×10^8^ intensity) by 120min while cells pre-treated with Cdt^WT^ (500ng/ml) reached maximum intensity (3.7×10^8^ intensity) at 150min. Also, when we employed the 96-well plate assay at 180min, macrophages pretreated with Cdt^WT^ (500ng/ml) exhibited a 40% decrease in DQ™-BSA fluorescence (Figure 6A and B) suggesting a reduction in phagolysosome formation. Cdt^R117A^ treatment did not alter DQ™-BSA fluorescence implicating phosphatase activity as the major contributor to decreased phagolysosome formation (Figure 6B). Lastly, neither Cdt^WT^ nor Cdt^R117A^, altered lysosome integrity (S.Figure 2)

**Figure 6.**
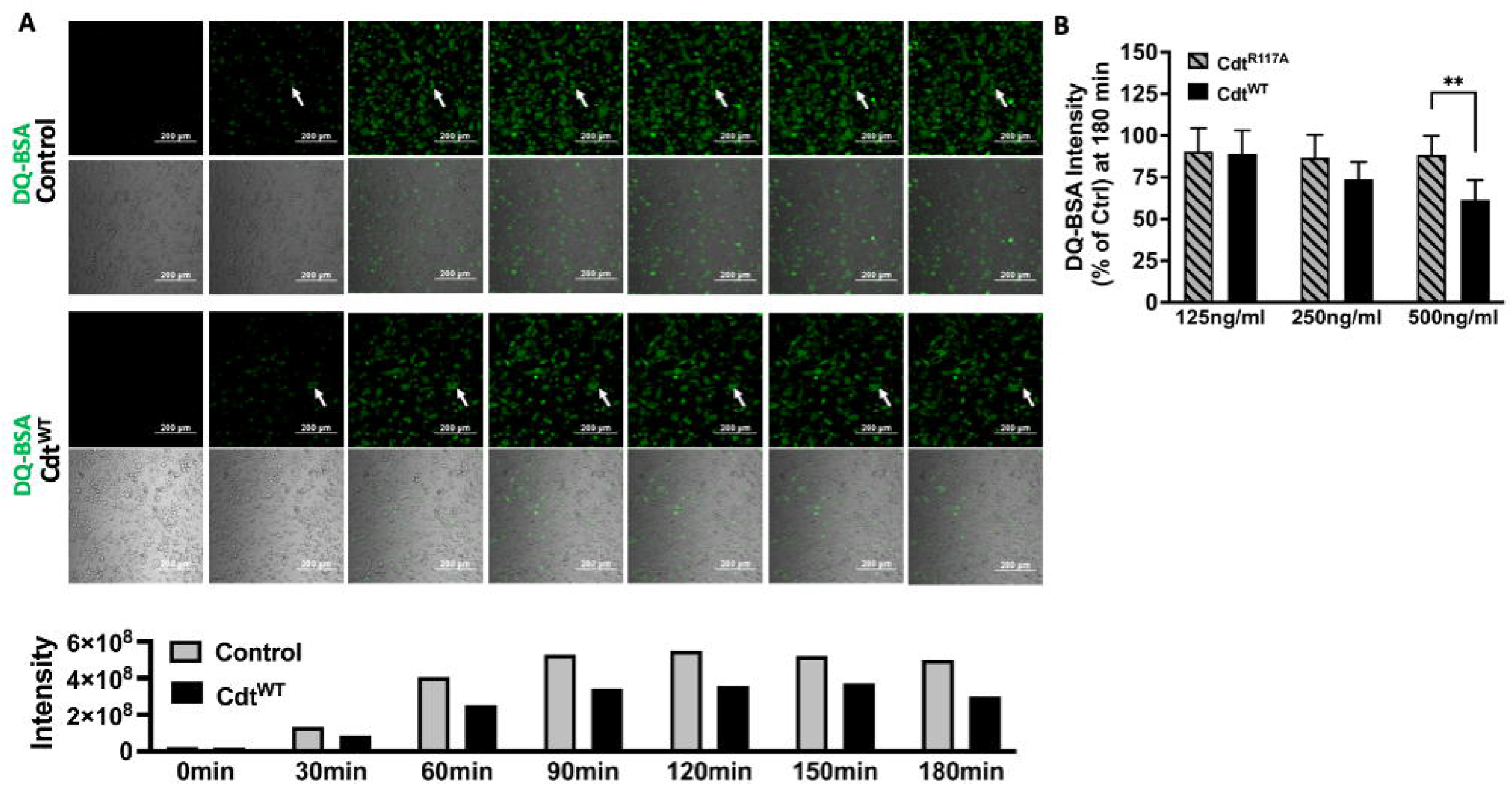
Cdt decreases phagosome maturation. A. Time lapse imaging of DQ™-BSA in THP1 macrophages pre-treated with Cdt^WT^ or control (untreated) for 4 hours. DQ™-BSA fluorescence is indicated by white arrows every 30 minutes over 3 hours (Top panel: Untreated, Bottom panel: Cdt^WT^). Intensity measurements are plotted over the same time course in the corresponding graph. B. DQ™-BSA intensity at 180 minutes, using 96 well plate format is shown as a function of increasing concentrations of Cdt^WT^ or Cdt^R117A^ as percent of control (untreated) as in Methods. Data represent mean +/- STDEV (5 wells per individual experiment with 1×10^6^ cells/well) and compared using Student’s t-test. **, p-value <0.005 Cdt^WT^ vs Cdt^R117A^.

The studies detailed above suggest that Cdt^WT^ decreases phagosome maturation by modulating PI pools. In this final series of experiments, we sought to determine if Cdt^WT^ alters macrophage bactericidal function by assessing *Aa* survival. When THP1 macrophages were inoculated with wild type *Aa* (D7S), or the mutant *Aa* lacking Cdt (D7SΔCdt) at a multiplicity of infection (MOI) of 1:10 for 14 hours, both bacterial strains exhibited virtually the same normalized CFUs (Figure 7A). In macrophages that were pretreated with Cdt^WT^ (500ng/ml) and subsequently inoculated with either wild type *Aa* or Cdt deficient mutant *Aa*, there was a 28% increase in *Aa* (D7S), and a 20% increase in *Aa* (D7SΔCdt) survival. Thus, pretreatment of THP1 derived macrophages with Cdt^WT^ resulted in an increase in *Aa* survival and a decrease in bactericidal effect (Figure 7A). When we utilized an MOI of 1:100 for 14 hours, macrophage cell death was observed (data not shown).

**Figure 7.**
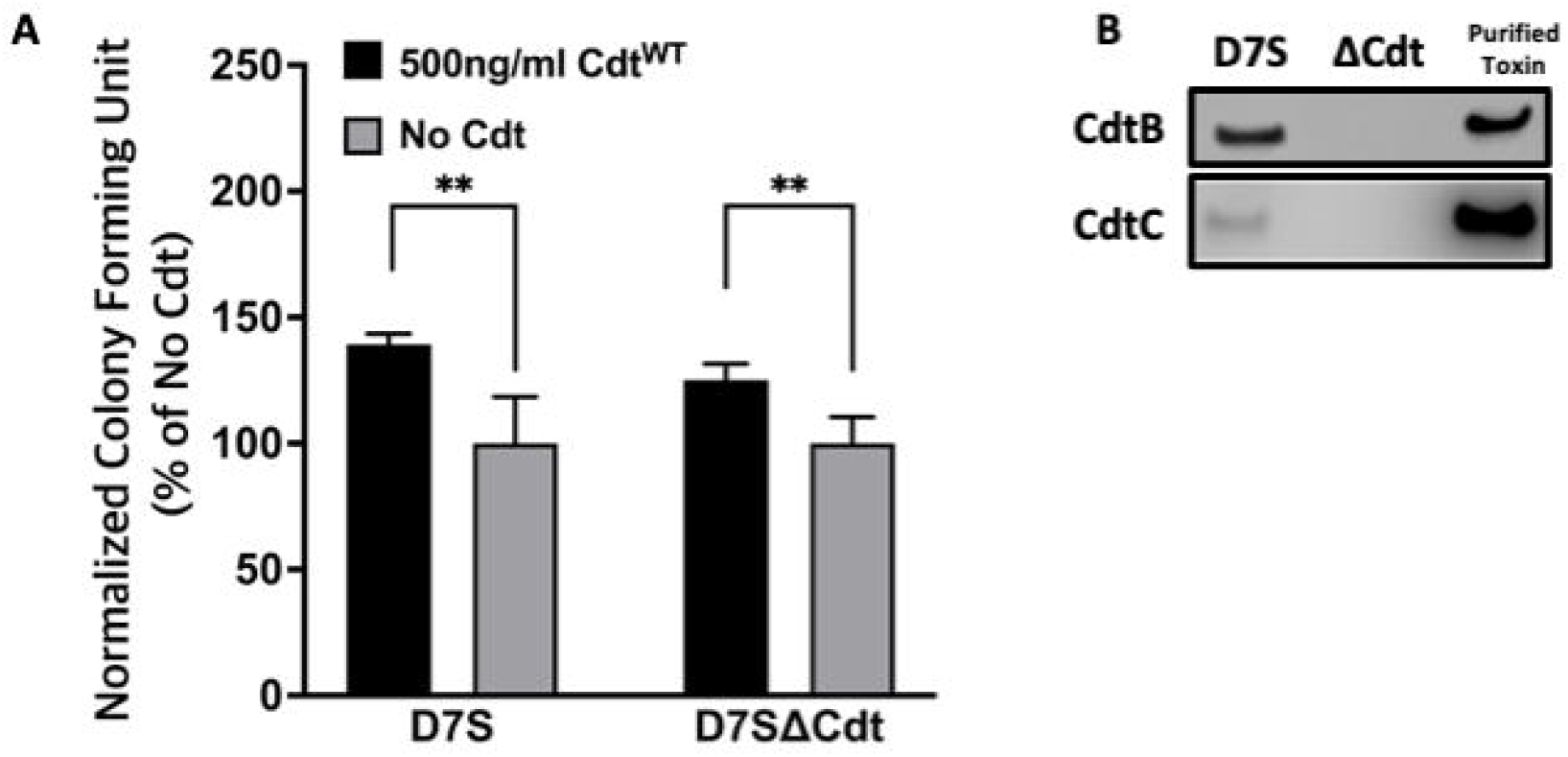
Cdt pretreatment enhances *Aa* survival. A. THP1 macrophages were pre-treated with 500ng/ml Cdt^WT^ for 4 hours or control (untreated). Cells were subsequently incubated with D7S and D7SΔCdt for 14hrs. Cells treated as in Methods and lysates were plated on agar plate and CFU units were determined. Data represent mean +/- STDEV (n=3) as percent of control (untreated) as described in Methods and compared using Student’s t-test **, p<0.05 vs No Cdt. B. Confirmation of Cdt production in D7S and loss of Cdt in D7SΔCdt strains. D7S and D7SΔCdt were plated on agar and then cultured by broth media. Western blot analysis was performed on bacterial lysates and positive control (purified Cdt) using mouse antibodies for Cdt subunits to confirm the mutations.

## DISCUSSION

In the oral cavity, *Aa* is a risk factor for gingival inflammation and localized aggressive periodontitis (Stage 3 or 4 and Grade C with molar-incisor pattern);we propose that Cdt contributes to the virulence properties of *Aa* [4, 6]. We have documented a novel intoxication profile for Cdt, as a tri-perditious toxin and shown that the manner of intoxication is cell type specific [13, 40]. Human monocytes and macrophages are resistant to *Aa-*Cdt induced apoptosis, while lymphocytes become apoptotic [40, 41]. In macrophages, we show Cdt, induces a robust pro-inflammatory response that involves activation of the NLRP3 inflammasome as well as the non-canonical inflammasome with cytokine release dependent on gasdermin cleavage [32].

Ando-Suguimoto, and colleagues suggested that the Cdt may modulate macrophage function in *Aa* infected sites by impairing phagocytosis using murine macrophages [38]. Our current studies expand on this observation to describe the molecular mechanism by which Cdt exerts its effect and contributes to *Aa* pathogenicity by creating an intracellular survival niche in human macrophages. We have shown that once internalized, the active CdtB subunit alters intracellular phosphoinositide pools, with the enzymatic depletion of PIP3 resulting in blockade of the PI-3K signaling pathway leading to both Akt inactivation and GSK3β activation [11]. In these studies, we explored the effect of Cdt mediated changes in phosphoinositide pools on the modulation of host phagocytic function. Intracellular phosphoinositide pools not only serve as signaling platforms but also mediate phagosome maturation and pathogen degradation [14-16]. Phagosomes are dynamic structures that interact with endosomes in a process involving acquisition and release of membrane and luminal components as the phagosome matures to a phago-lysosome [14-16]. This maturation process is controlled by recruitment of proteins, such as Rab5 and Rab7 (and other effector proteins), which is regulated by PI distribution [18, 19]. Specific phosphoinositide species act as zip-codes ensuring that the maturing phagosome is targeted towards lysosomes for fusion and degradation as illustrated schematically in Figure 8. PI(3,4,5)P3 and PI(4,5)P2 play critical roles in phagosome formation and PI3P and PI(3,4)P2 in phagosome maturation.

**Figure 8.**
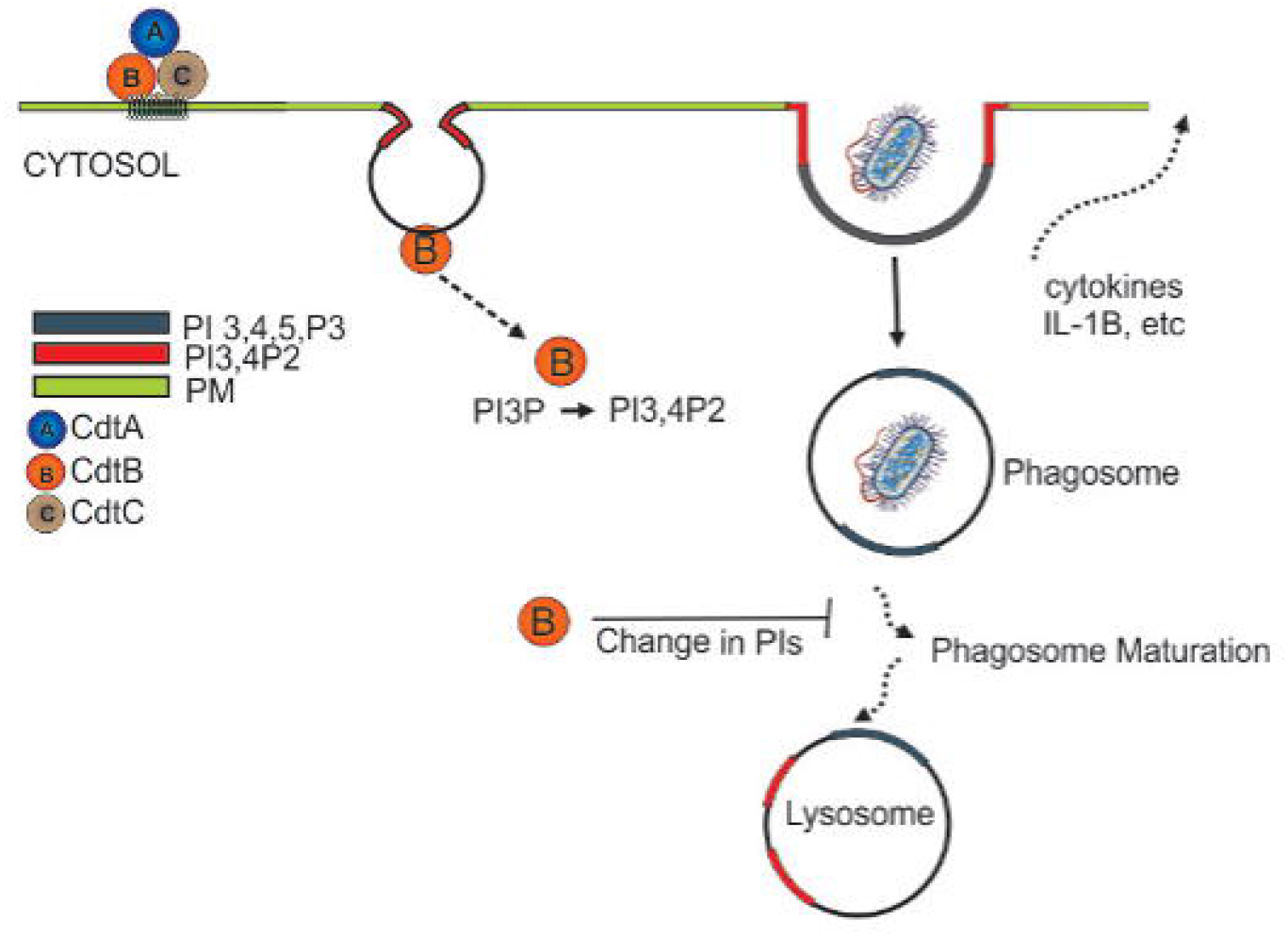
Overview of Cdt internalization and the effect on the phagosome. After the internalization, CdtB act as a phosphatase and convert the PI(3,4,5)P3 to PI(3,4)P2. Change in the PI pool leads to manipulation of phagosome maturation.

Herein we show that Cdt^WT^, as expected stimulates a decrease in PI(3,4,5)P3 and a corresponding increase in PI(3,4)P2 which is localized to distinct domains and surrounds phagocytosed latex beads (Figs. 1 and 3). The extent of PI(3,4)P2 association with latex beads is clearly visible in the presence of Cdt^WT^ but is undetectable in cells treated with Cdt^R117A^ or in untreated cells (Figure 3). Importantly, PI(4,5)P2 levels remained unaltered (Figure 1) and pHrodo™ (Figure 2) and DQ™-BSA (Figure 6) uptake did not vary between Cdt^WT^ and Cdt^R117A^ suggesting that the initial step in uptake, a PI4,5,P2 dependent process that recruits actin-modifying proteins is intact. Our Rab5 and EEA1 latex bead association studies suggest that Cdt^WT^ phosphatase activity modulates a step downstream of uptake in the maturation process. EEA1 recruitment to latex beads was not sensitive to Cdt^WT^ treatment (Figure 5B). In contrast association of Rab5, an early phagosome/endosome associated GTPase with latex beads decreased upon Cdt^WT^ treatment Rab5-latex bead association was decreased (Figure 5A). Whether this decrease is due to alteration in the levels of PI3P, is currently under investigation. Early Rab5 positive endosomes/phagosomes subsequently become late Rab7 positive structures [18, 42]. Thus, diminished Rab5 phagosome association can be indicative of stalled phagosome or slowed phagosome maturation. In fact, Cdt^WT^ treatment resulted in decreased phagosome maturation and fusion with lysosomes as indicated in Figure 6. Collectively, these studies provide a molecular mechanism to explain diminished phagocytic capacity observed in murine macrophages upon *Aa*Cdt treatment [38].

These studies provide molecular insight into how *Aa*Cdt subverts phagocytosis and creates a survival niche for subsequent infections (Figure 7). *Aa*, specifically due to its CdtB phosphatase activity, can be added to the rapidly growing list of pathogens that have evolved to evade antimicrobial effects of macrophages thus crippling the effective immune response [43]. In the context of periodontal disease, this effect of *Aa*Cdt provides a molecular mechanism by which *Aa* functions in the oral cavity to provide a “survival” niche in the gingival crevice for other pathogenic microorganisms which collectively contribute to LAP pathogenesis [3, 6, 44]. *Aa* is present in healthy oral flora, however the transition from health to disease is reflected in the habitat within which *Aa* exists [6]. Under healthy conditions, *Aa* dwell and colonize above the gum-line. In the disease state, *Aa* transits below the gum-line, to the subgingival niche, an area with less oxygen [6]. This localization favors *Aa* as it is a facultative anaerobic gram-negative bacterium. *Aa* utilize three major toxins. apiA gives adhesion, invasion, and resistance to the complement system [6]. Leukotoxin causes leukocyte apoptosis [5, 45]. Cdt causes cell cycle arrest, apoptosis in T cells and epithelial cells, upregulates the pro-inflammatory cytokines in macrophages [13, 40, 41], and with current study, it also decreased the phagocytic effect of macrophage. We predict that these disruptions could impair the body’s defense system by decreasing the antimicrobial effect, causes dysbiosis, and exacerbate the inflammatory response. These combined effects of *Aa* strongly suggest that it may be an initiator of LAP thereby preparing the niche for other pathogens. This interpretation can also be applicable to other Cdt producing pathogen as a new mechanism of hijacking host function to create pathologically favorable condition.

Lastly, Our observation that CdtB phosphatase activity slows phagosome maturation likely through a Rab 5 effector provides a molecular mechanism for the observation that *Campylobacter jejuni*, Cdt show elevated toxicity in cells over expressing Rab5 [46]. We predict this is due to CdtB-mediated decreased phagosome maturation hence increased *Campylobacter jejuni* survival and this prediction remains to be tested.

## Acknowledgements

these studies were supported by NIDCR: DE023071-07 (BJS and KBB) and 1K08DE032119-01 (TJK). The authors thank Dr. Casey Chen from University Southern California Herman Ostrow School of Dentistry for providing *Aa* D7S-SA (wild type *Aa*) and D7S-SA CHE001 (Cdt deficient *Aa* mutant) and Dr. Anuradha Dhingra for her assistance with live-cell imaging and the PDM live cell imaging core.

## SUPPLEMTENTAL FIGURES

**Figure S1.**
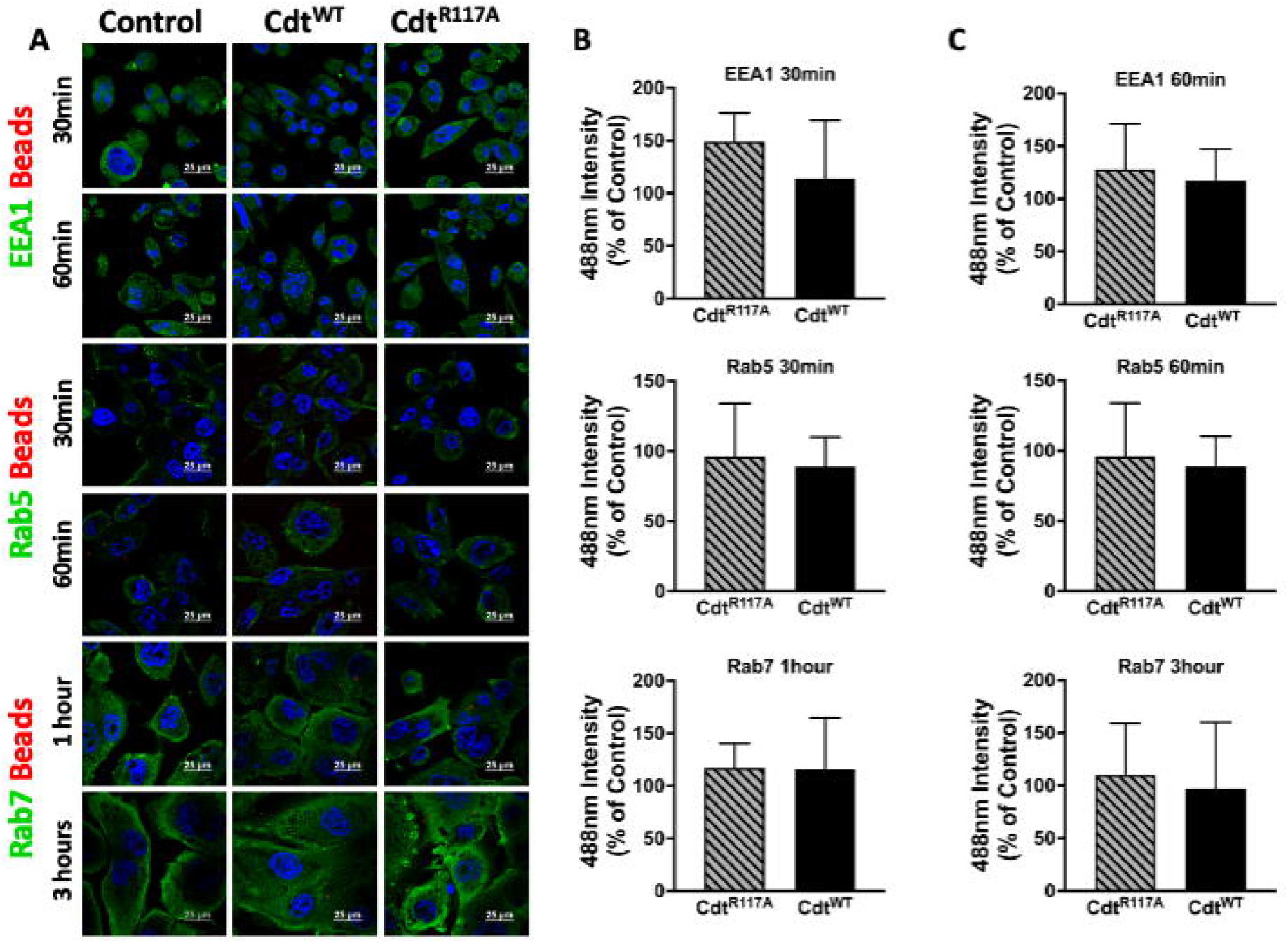
Effector proteins, Rab5, EEA1, and Rab7 levels were unaltered upon Cdt treatment. THP1 macrophages treated for 4 hours with Cdt^WT^, Cdt^R117A^, or control (untreated) prior to synchronized uptake of opsonized beads were fixed (at 30min and 60min for Rab5 and EEA1, at 1hr and 3hr for Rab7) and stained for Rab5, EEA1, and Rab7 as in Methods. Data represent mean +/- STDEV of protein intensity (5 different fields with about 12 cells/field) as percent to control and compared using Student’s t-test.

**Figure S2.**
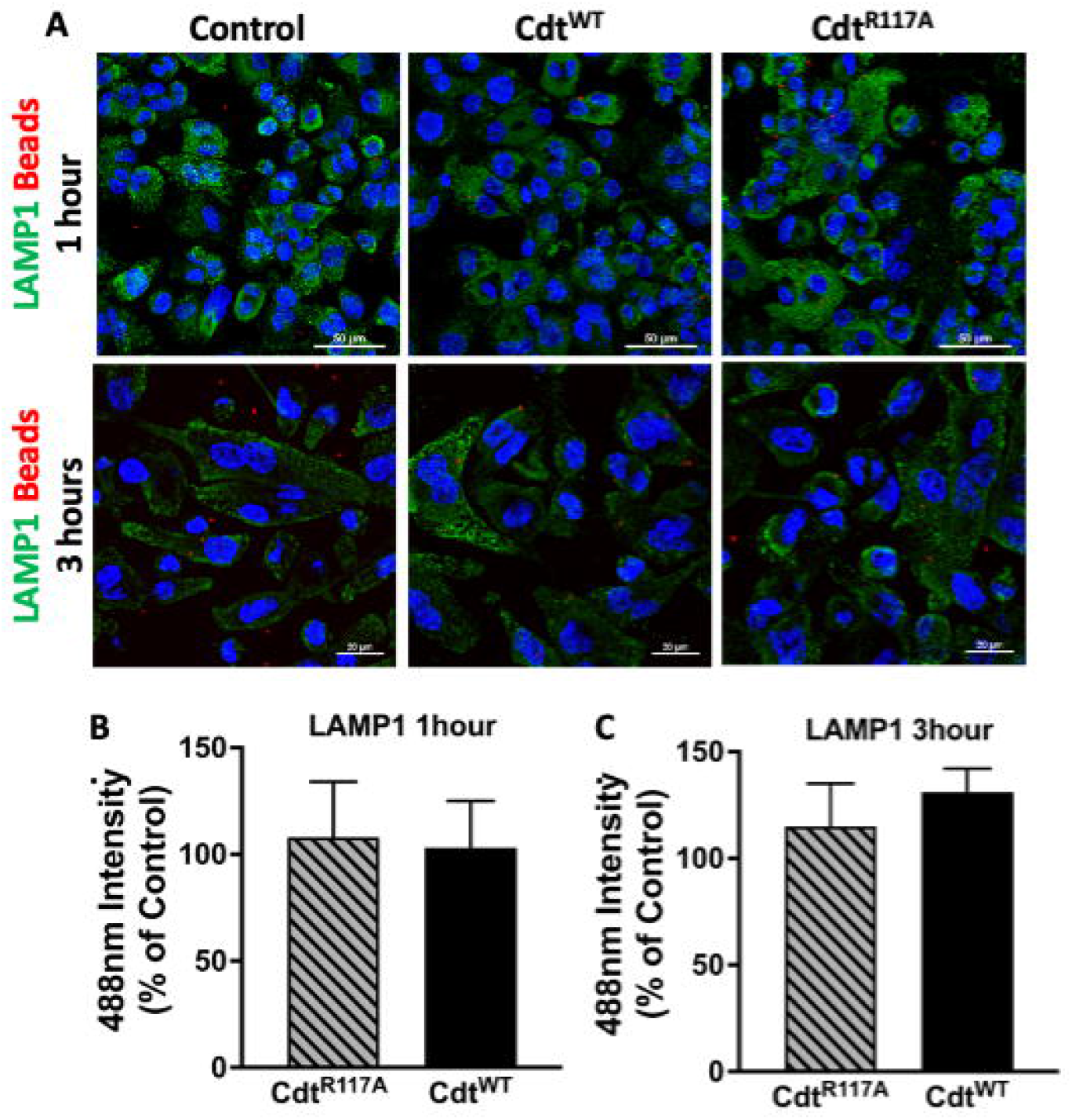
Cdt treatment does not alter LAMP1. A. LAMP1 staining in THP1 macrophages pre-treated with Cdt^WT^ (500ng/ml), Cdt^R117A^ or control (untreated) prior to the addition of opsonized beads as in Methods. B. LAMP1 fluorescence intensity at 1 hour (Panel B) or 3 hours (Panel C). Data represent mean +/- STDEV of LAMP1 intensity (5 fields per condition with average of 12 cells/field) as a percent to control and compared by using Student’s t-test.

## REFERENCE

1. Hienz, S.A., S. Paliwal, and S. Ivanovski, Mechanisms of Bone Resorption in Periodontitis. J Immunol Res, 2015. 2015: p. 615486.

2. Kornman, K.S., Mapping the pathogenesis of periodontitis: a new look. J Periodontol, 2008. 79(8 Suppl): p. 1560–8.

3. Teughels, W., et al., Treatment of aggressive periodontitis. Periodontol 2000, 2014. 65(1): p. 107–33.

4. Tonetti, M.S., H. Greenwell, and K.S. Kornman, Staging and grading of periodontitis: Framework and proposal of a new classification and case definition. J Periodontol, 2018. 89 Suppl 1: p. S159–S172.

5. Oscarsson, J., et al., Tools of Aggregatibacter actinomycetemcomitans to Evade the Host Response. J Clin Med, 2019. 8(7).

6. Fine, D.H., A.G. Patil, and S.K. Velusamy, Aggregatibacter actinomycetemcomitans (Aa) Under the Radar: Myths and Misunderstandings of Aa and Its Role in Aggressive Periodontitis. Front Immunol, 2019. 10: p. 728.

7. Fine, D.H., et al., Aggregatibacter actinomycetemcomitans as an early colonizer of oral tissues: epithelium as a reservoir? J Clin Microbiol, 2010. 48(12): p. 4464–73.

8. Johnson, W.M. and H. Lior, A new heat-labile cytolethal distending toxin (CLDT) produced by Campylobacter spp. Microb Pathog, 1988. 4(2): p. 115–26.

9. Johnson, W.M. and H. Lior, A new heat-labile cytolethal distending toxin (CLDT) produced by Escherichia coli isolates from clinical material. Microb Pathog, 1988. 4(2): p. 103–13.

10. Huang, G., et al., The Active Subunit of the Cytolethal Distending Toxin, CdtB, Derived From Both Haemophilus ducreyi and Campylobacter jejuni Exhibits Potent Phosphatidylinositol-3,4,5-Triphosphate Phosphatase Activity. Front Cell Infect Microbiol, 2021. 11: p. 664221.

11. Shenker, B.J., et al., The toxicity of the Aggregatibacter actinomycetemcomitans cytolethal distending toxin correlates with its phosphatidylinositol-3,4,5-triphosphate phosphatase activity. Cell Microbiol, 2016. 18(2): p. 223–43.

12. Boesze-Battaglia, K., et al., A Journey of Cytolethal Distending Toxins through Cell Membranes. Front Cell Infect Microbiol, 2016. 6: p. 81.

13. Shenker, B.J., et al., Blockade of the PI-3K signalling pathway by the Aggregatibacter actinomycetemcomitans cytolethal distending toxin induces macrophages to synthesize and secrete pro-inflammatory cytokines. Cell Microbiol, 2014. 16(9): p. 1391–404.

14. Jeschke, A. and A. Haas, Sequential actions of phosphatidylinositol phosphates regulate phagosome-lysosome fusion. Mol Biol Cell, 2018. 29(4): p. 452–465.

15. Jeschke, A., et al., Phosphatidylinositol 4-phosphate and phosphatidylinositol 3-phosphate regulate phagolysosome biogenesis. Proc Natl Acad Sci U S A, 2015. 112(15): p. 4636–41.

16. Wallroth, A. and V. Haucke, Phosphoinositide conversion in endocytosis and the endolysosomal system. J Biol Chem, 2018. 293(5): p. 1526–1535.

17. Fratti, R.A., et al., Interdependent assembly of specific regulatory lipids and membrane fusion proteins into the vertex ring domain of docked vacuoles. J Cell Biol, 2004. 167(6): p. 1087–98.

18. Vieira, O.V., et al., Modulation of Rab5 and Rab7 recruitment to phagosomes by phosphatidylinositol 3-kinase. Mol Cell Biol, 2003. 23(7): p. 2501–14.

19. Shin, H.W., et al., An enzymatic cascade of Rab5 effectors regulates phosphoinositide turnover in the endocytic pathway. J Cell Biol, 2005. 170(4): p. 607–18.

20. Yarwood, R., et al., Membrane trafficking in health and disease. Dis Model Mech, 2020. 13(4).

21. Vicinanza, M., et al., Phosphoinositides as regulators of membrane trafficking in health and disease. Cell Mol Life Sci, 2008. 65(18): p. 2833–41.

22. Shenker, B.J., et al., Actinobacillus actinomycetemcomitans cytolethal distending toxin (Cdt): evidence that the holotoxin is composed of three subunits: CdtA, CdtB, and CdtC. J Immunol, 2004. 172(1): p. 410–7.

23. Shenker, B.J., et al., Expression of the cytolethal distending toxin (Cdt) operon in Actinobacillus actinomycetemcomitans: evidence that the CdtB protein is responsible for G2 arrest of the cell cycle in human T cells. J Immunol, 2000. 165(5): p. 2612–8.

24. Lally, E.T., et al., Aggregatibacter actinomycetemcomitans LtxA Hijacks Endocytic Trafficking Pathways in Human Lymphocytes. Pathogens, 2020. 9(2).

25. Boesze-Battaglia, K., et al., Internalization and Intoxication of Human Macrophages by the Active Subunit of the Aggregatibacter actinomycetemcomitans Cytolethal Distending Toxin Is Dependent Upon Cellugyrin (Synaptogyrin-2). Front Immunol, 2020. 11: p. 1262.

26. Boesze-Battaglia, K., et al., Internalization of the Active Subunit of the Aggregatibacter actinomycetemcomitans Cytolethal Distending Toxin Is Dependent upon Cellugyrin (Synaptogyrin 2), a Host Cell Non-Neuronal Paralog of the Synaptic Vesicle Protein, Synaptogyrin 1. Front Cell Infect Microbiol, 2017. 7: p. 469.

27. Reyes-Reveles, J., et al., Phagocytosis-dependent ketogenesis in retinal pigment epithelium. J Biol Chem, 2017. 292(19): p. 8038–8047.

28. Segawa, T., et al., Inpp5e increases the Rab5 association and phosphatidylinositol 3-phosphate accumulation at the phagosome through an interaction with Rab20. Biochem J, 2014. 464(3): p. 365–75.

29. Kissing, S., et al., Vacuolar ATPase in phagosome-lysosome fusion. J Biol Chem, 2015. 290(22): p. 14166–80.

30. Magenau, A., et al., Phagocytosis of IgG-coated polystyrene beads by macrophages induces and requires high membrane order. Traffic, 2011. 12(12): p. 1730–43.

31. Kapellos, T.S., et al., A novel real time imaging platform to quantify macrophage phagocytosis. Biochem Pharmacol, 2016. 116: p. 107–19.

32. Shenker, B.J., et al., Cytolethal distending toxin-induced release of interleukin-1beta by human macrophages is dependent upon activation of glycogen synthase kinase 3beta, spleen tyrosine kinase (Syk) and the noncanonical inflammasome. Cell Microbiol, 2020. 22(7): p. e13194.

33. Frost, L.S., et al., The Contribution of Melanoregulin to Microtubule-Associated Protein 1 Light Chain 3 (LC3) Associated Phagocytosis in Retinal Pigment Epithelium. Mol Neurobiol, 2015. 52(3): p. 1135–1151.

34. Frost, L.S., et al., The Use of DQ-BSA to Monitor the Turnover of Autophagy-Associated Cargo. Methods Enzymol, 2017. 587: p. 43–54.

35. Nalbant, A., et al., Induction of T-cell apoptosis by Actinobacillus actinomycetemcomitans mutants with deletion of ltxA and cdtABC genes: possible activity of GroEL-like molecule. Oral Microbiol Immunol, 2003. 18(6): p. 339–49.

36. Sreenivasan, P.K., D.H. Meyer, and P.M. Fives-Taylor, Factors influencing the growth and viability of Actinobacillus actinomycetemcomitans. Oral Microbiol Immunol, 1993. 8(6): p. 361–9.

37. Sharma, A. and A. Puhar, Gentamicin Protection Assay to Determine the Number of Intracellular Bacteria during Infection of Human TC7 Intestinal Epithelial Cells by Shigella flexneri. Bio Protoc, 2019. 9(13): p. e3292.

38. Ando-Suguimoto, E.S., et al., The cytolethal distending toxin of Aggregatibacter actinomycetemcomitans inhibits macrophage phagocytosis and subverts cytokine production. Cytokine, 2014. 66(1): p. 46–53.

39. Perez-Stuardo, D., et al., Non-lysosomal Activation in Macrophages of Atlantic Salmon (Salmo salar) After Infection With Piscirickettsia salmonis. Front Immunol, 2019. 10: p. 434.

40. Scuron, M.D., et al., The Cytolethal Distending Toxin Contributes to Microbial Virulence and Disease Pathogenesis by Acting As a Tri-Perditious Toxin. Front Cell Infect Microbiol, 2016. 6: p. 168.

41. Shenker, B.J., et al., Induction of cell cycle arrest in lymphocytes by Actinobacillus actinomycetemcomitans cytolethal distending toxin requires three subunits for maximum activity. J Immunol, 2005. 174(4): p. 2228–34.

42. Mottola, G., The complexity of Rab5 to Rab7 transition guarantees specificity of pathogen subversion mechanisms. Front Cell Infect Microbiol, 2014. 4: p. 180.

43. Sarantis, H. and S. Grinstein, Subversion of phagocytosis for pathogen survival. Cell Host Microbe, 2012. 12(4): p. 419–31.

44. Loe, H. and L.J. Brown, Early onset periodontitis in the United States of America. J Periodontol, 1991. 62(10): p. 608–16.

45. Korostoff, J., et al., Actinobacillus actinomycetemcomitans leukotoxin induces apoptosis in HL-60 cells. Infect Immun, 1998. 66(9): p. 4474–83.

46. Chen, M.X., et al., Rab5a Promotes Cytolethal Distending Toxin B-Induced Cytotoxicity and Inflammation. Infect Immun, 2020. 88(10).

